# EVODEX: A Mechanistic Framework for Extracting, Structuring, and Predicting Enzymatic Reactivity

**DOI:** 10.1101/2025.06.19.660615

**Authors:** Lais Lastre Conceicao, Haohong Lin, Doris Tai, Gongao Xue, Han Zhang, Chenyan Zhang, J. Christopher Anderson

## Abstract

Accurately modeling enzymatic reactivity is essential for synthesis planning, mass spectrometry interpretation, and biochemical knowledge extraction, but existing reaction datasets are often incomplete or inconsistently annotated. EVODEX introduces two abstractions to address this: partial reactions, which isolate atom-mapped substrate-product transformations from complex biochemical reactions, and Electronic Reaction Operators (EROs), which encode the bonding and orbital environment of atoms undergoing change. From 349,458 curated reactions in EnzymeMap, EVODEX constructs over 186,000 partial reactions and mines a core set of 1,404 EROs, each supported by at least 10 examples. Together, these explain 91 percent of test reactions at the formula level and 62 percent at the mechanistic level. A further filtered synthesis subset of 436 one-to-one operators that exclude ubiquitous metabolites provides a compact and generalizable vocabulary of single-step enzymatic transformations. EVODEX operators are auditable, reusable, and available as part of an open-source Python package with datasets, Jupyter notebooks, and a browsable website.

## Introduction

Predicting biochemical reactivity is a foundational task in computational biochemistry, central to applications such as synthesis planning, mass spectrometry interpretation, and pathway reconstruction. This task depends on understanding two distinct but interrelated phenomena: the atomic transformations dictated by bonding and electron movement, and the enzymatic specificity that governs substrate orientation and activation. EVODEX (Enzymatic Validation Operator Differentiation and Extraction) addresses the first of these by providing a mechanistic framework for analyzing enzymatic transformations through precise structural abstractions.

At the core of EVODEX is the Electronic Reaction Operator (EVODEX-E), a deterministic representation of a reaction’s electronic mechanism. These operators capture the sigma and pi bonding environment around reactive atoms, encoding molecular orbital context in a format consistent with traditional arrow-pushing conventions. Each operator is derived directly from an atom-mapped reaction, extracted without manual rule design, and interpretable by both chemists and software.

A range of prior tools have defined transformation rules to support pathway generation, metabolite prediction, and retrosynthesis. BNICE introduced generalized transformation rules that approximate electron movement, though without a formal bonding or orbital framework, and its rule sets are not publicly available^6,7^. RetroRules^8^ extends this approach using a published dataset of automatically extracted rules defined by fixed-radius neighborhoods around reactive centers, with radius-1 rules closely matching EVODEX’s nearest-neighbor abstractions. Other systems such as SimPheny^9^ and MINE 2.0^10^ use manually curated rule libraries derived from known reactions, while BioTransformer^11,12^ and enviPath^13^ apply small, expert-defined sets of transformations aimed at predicting drug metabolism or annotating mass spectra. A recent study^14^ by García-Jiménez et al. curated a compact list of tailoring transformations. They applied these to predicted polyketide and nonribosomal peptide scaffolds in order to model biosynthetic elaborations. These systems typically rely on fully balanced reactions and assume predefined cofactors, which limits their coverage of real-world biochemical data. EVODEX overcomes this constraint by working with partial reactions that preserve mapped atomic changes even when cofactors or side products are unknown. This does not require balanced equations, predefined cofactors, or manual rule curation.

Working with partial reactions introduces significant technical challenges. These reactions are typically chemically unbalanced, requiring explicit hydrogen accounting to preserve valency. Most cheminformatics libraries, including RDKit^15^, ChemAxon^16^, and Indigo^17^, treat hydrogens as special cases, making consistent representation difficult.

Additionally, partial reactions typically contain unmapped atoms that appear on only one side, raising ambiguities that must be resolved through well-defined rules. To manage these issues, EVODEX defines a set of operator types. These include EVODEX-E, -C, -N, and their matched variants, that capture bonding, mapping, and structural scope at multiple levels of abstraction.

EVODEX.1 formalizes this system in a structured and deployable form. It includes curated datasets of ten operator types, a pip-installable Python package, Jupyter notebook demonstrations, and a static website for browsing all operators and their source reactions. These tools support three core applications. In synthesis, EVODEX-E operators generate plausible products from known substrates. In mass spectrometry, EVODEX-M operators explain observed mass differences by linking them to chemical transformations. In reaction plausibility evaluation, EVODEX-C and EVODEX-F operators assess whether proposed transformations are chemically and mechanistically consistent with known patterns. Figure 1 illustrates these use cases.

**Figure 1.**
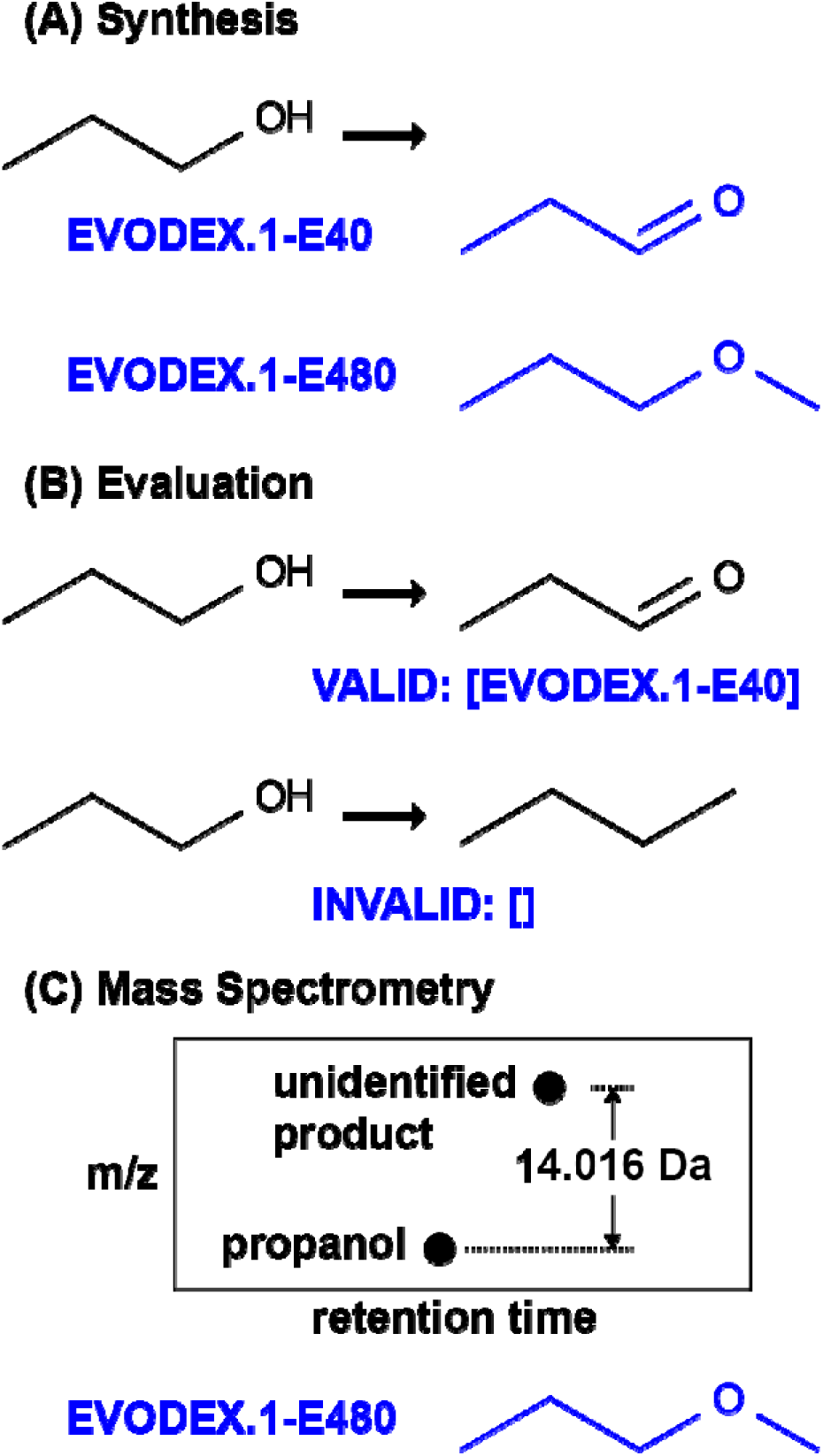
Illustration of EVODEX applications. (A) Provided a substrate structure, EVODEX operators predict mechanistically-plausible products that can be reached in one enzymatic step without consideration of specificity. (B) A proposed reaction is evaluated for mechanistic plausibility by matching to mined EVODEX operators. (C) Observed ion mass differences in mass spectrometry data can be interpreted by EVODEX to predict plausible products and the mechanism of their formation. See notebook examples^3,4,5^ for full results; partial output shown in blue.

## Methods

The EVODEX software suite is publicly available through the main GitHub repository^1^ under the open-source MIT License. This section describes the construction of EVODEX.1, the current release of the EVODEX framework. It consists of two main components: the evodex Python package, which contains the core cheminformatics algorithms for reaction operator extraction and manipulation, and the pipeline module, which orchestrates the full operator mining workflow. This spans all stages from raw data processing to the generation of final operator tables and a static HTML website. The EVODEX.1 website^2^ presents browsable operator pages with visualizations and metadata for each entry. All resulting CSV datasets are bundled within the repository and included in the PyPI installable package. Users can access the complete system without building from source or running the pipeline manually. Example Jupyter notebooks are provided both in the repository and online^3,4,5^, illustrating key use cases such as operator projection, mass shift analysis, and product prediction.

### Mining Evodex

Although EVODEX operators are defined through deterministic and independently computable algorithms, the practical task of mining them from biochemical datasets is not straightforward. EnzymeMap^18^. was selected as the foundation due to its curated and high-accuracy mappings. However, several real-world challenges, such as the scale of the dataset, the need for explicit hydrogen reconstruction, and the presence of incorrect mappings, made the process technically intensive. Mining required extensive conversion, hashing, validation, and pruning, each of which introduced potential points of failure and demanded substantial preprocessing to produce a reliable and consistent set of operators. Figure 2 summarizes this workflow, from raw reactions to EVODEX-R, -P, and -F abstractions, including a synthesis-focused subset.

**Figure 2:**
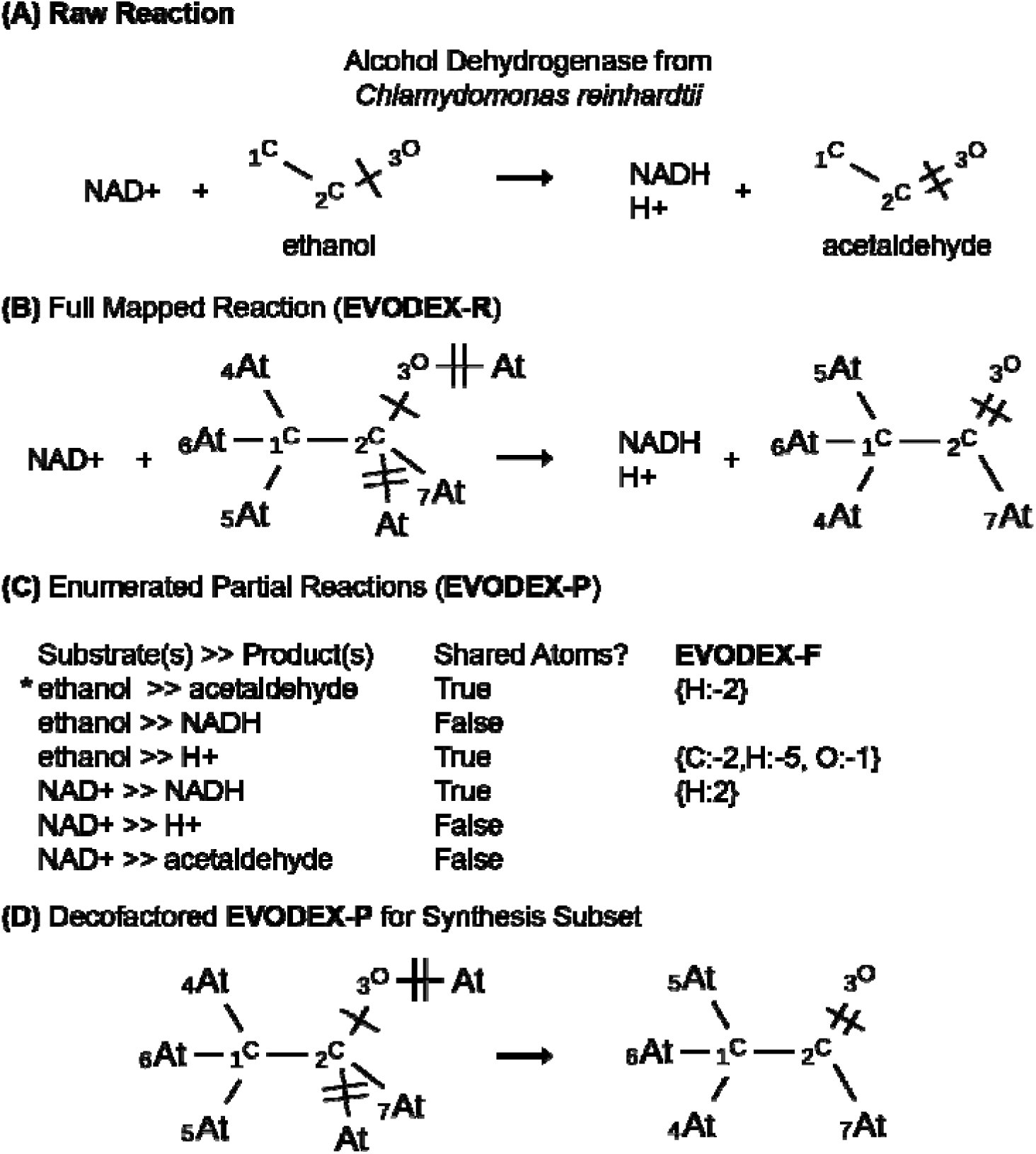
Pipeline for Mining. (A) The raw input contains complete reactions, including the specific enzyme, and atom maps on heavy atoms. (B) The enzyme is omitted, the Hydrogens are converted to Astatine and mapped to generate EVODEX-R abstractions. (C) All partial reaction permutations for each EVODEX-R are computed and evaluated for each molecule contributing shared atoms. Valid ones become EVODEX-P and further abstracted to EVODEX-F, M, and reaction operators. (D) One of the EVODEX-P enumerations, indicated with an asterisk, also results from decofactoring wherein all of a list of ubiquitous metabolites are removed. The resulting subset is used by synthesis algorithms as a more focused set that eliminates unnecessary considerations on the availability of common metabolites.

The operator mining workflow is divided into seven distinct phases, each responsible for a specific part of the EVODEX.1 construction process. Each stage of the pipeline is implemented as a separate module, allowing for independent execution and debugging. The pipeline can be orchestrated by a top-level script, run_pipeline.py, which sequentially executes each stage. The entire pipeline was developed and executed in a Python environment using Visual Studio on a Mac mini (Apple M1, 8-core CPU, 8-core GPU, 16GB RAM). The execution time for the full pipeline is 3.6 hours. Details on building and running the pipeline are available on the GitHub repo^1^.

### Operator Extraction

The core component of the EVODEX pipeline is the extract_operator method in operators.py, which inputs a mapped reaction and outputs a reaction operator. All reaction operator indices (E, Em, C, Cm, N, Nm) are generated from this method using different combinations of Boolean arguments. These include: include_stereochemistry (whether to preserve stereochemical labels), include_sigma (whether to include atoms directly sigma-bonded to the reaction center), include_pi (whether to include atoms in conjugated pi systems adjacent to the center), include_unmapped_hydrogens (whether to include hydrogens that appear only on one side of the reaction), include_unmapped_heavy_atoms (similar, but for non-hydrogen atoms), and include_static_hydrogens (whether to retain non-reactive hydrogens consistently bonded to mapped atoms). Based on these flags, atoms are selected for inclusion in the operator. A duplicate of the original reaction is then pruned to contain only these atoms, and the result is returned as a SMIRKS string. This design is illustrated in Figure 3, which shows how different abstraction levels and inclusion choices (e.g., static hydrogens, conjugated atoms) affect the representation of a glutathione-catalyzed Michael addition.

**Figure 3:**
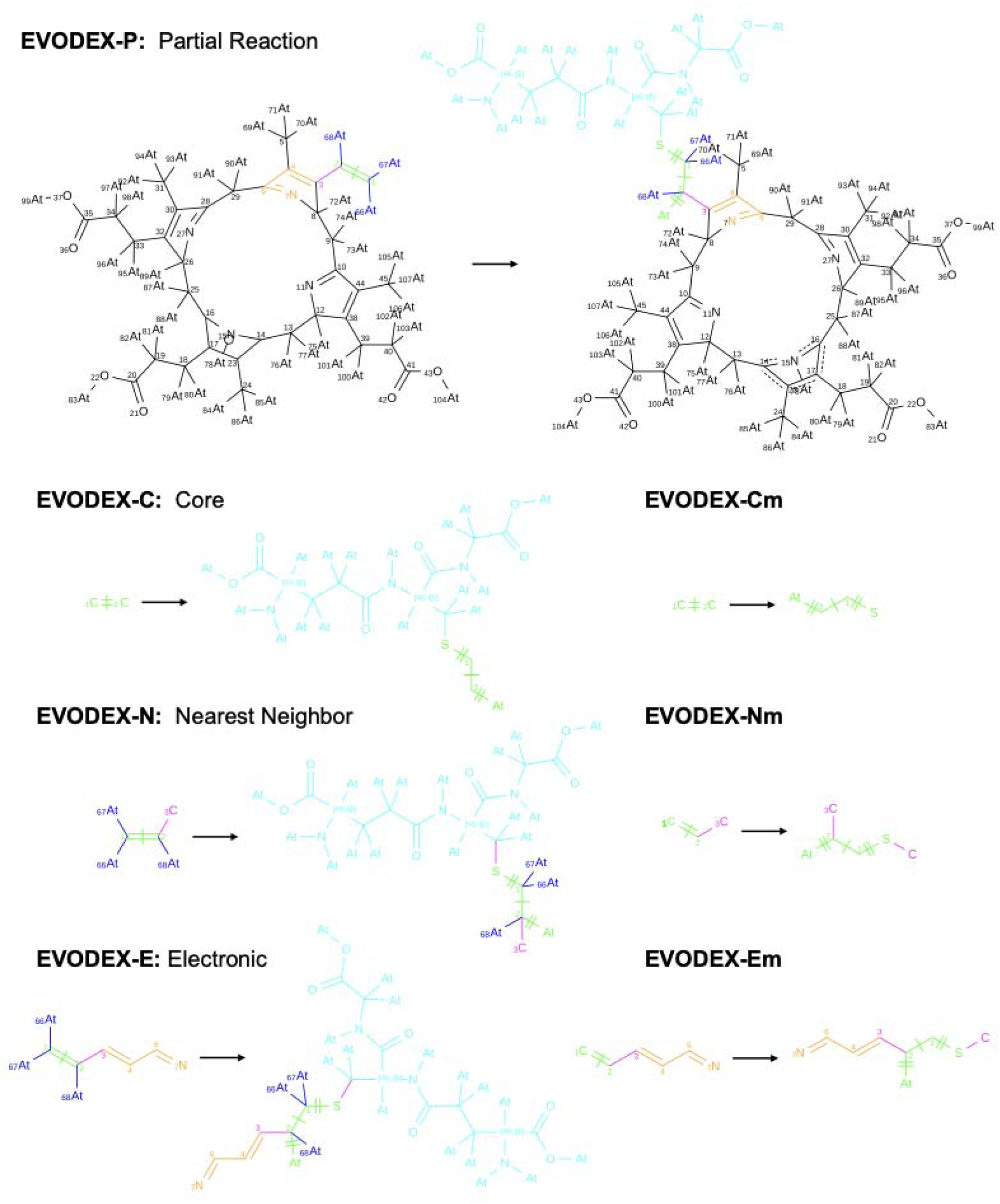
The glutathionylation of harderoporphyrinogen is shown as an EVODEX-P partial reaction, with glutathione omitted. The thiol group attacks an extended enamine in the substrate. Atom colors indicate role: green (core), magenta (sigma-bound), orange (pi-conjugated), black (mapped but unchanged), cyan (unmapped), and blue (static hydrogens). Operators are generated by abstraction type: Core (C), Nearest Neighbor (N), or Electronic (E), with matched variants (Cm, Nm, Em). Only the electronic abstraction captures the full reactive context in this case.

Operators are computed using Boolean flags that define variants such as C, N, E, and their matched counterparts (Cm, Nm, Em). The operators computed in the pipeline represent a carefully chosen subset of the broader abstraction space enabled by this framework. Complete operators (C, N, E) retain all mapped and unmapped atoms and are essential for product prediction, where atom imbalance across partial reactions must be resolved, while matched operators (Cm, Nm, Em) exclude unmapped atoms to isolate the minimal mechanistic transformation. The distinctions between Core (C), Nearest Neighbor (N), and Electronic (E) operators reflect increasing contextual scope— from the reactive center to sigma-bonded neighbors to extended conjugated systems.

While many operator variants are possible, not all are useful in practice. For example, omitting stereochemistry from complete operators can lead to ambiguous representations, such as flattening the chiral centers in glucose during glucosidation reactions. Conversely, retaining stereochemistry may introduce irrelevant distinctions such as differentiating R and S alcohols in oxidations where chirality has no mechanistic role. In such cases, more general operators derived from achiral analogs often exist and are favored automatically during dominance pruning, yielding concise and mechanistically appropriate forms. Figure 3 illustrates how abstraction scope affects operator construction: core and neighbor operators omit key features, while only the electronic operator captures the full conjugated system required to represent the reactive enamine in this glutathionylation.

### Reaction hashing

Custom hashing and consolidation procedures were developed because standard cheminformatics tools failed to generate consistent, representation-invariant identifiers for SMIRKS reactions. RDKit’s InChI and SMILES-based methods were sensitive to atom mapping, bond order, and kekulization. To address this, we used hybridization-encoded atomic labeling and canonical SMILES to define identity in terms of atom connectivity and local electronic context, rather than superficial structural form. Each molecule was reduced to a simplified form retaining only non-hydrogen, non-astatine atoms with all bonds set to single. Carbon atoms were labeled via isotopes to reflect their hybridization state, distinguishing saturated from oxidized environments. Substrates and products were hashed independently, then combined into a deterministic string (‘sorted_substrate_hashes >> sorted_product_hashes’) and SHA-256 hashed to produce the final operator identity. This ensures operators differing only by atom ordering, mapping, or resonance are grouped, while chemically distinct transformations remain separate.

### Phase 1: Data Preparation

The dataset used for mining EVODEX.1 was derived from the EnzymeMap^18^, itself based on curated biocatalysis reactions from BRENDA^19^. While EnzymeMap provides atom-mapped reactions, only heavy atoms are included. We augmented this data by adding hydrogens and then substituting them with astatines as placeholders, due to RDKit’s inconsistent behavior with explicit hydrogens. The substituted astatine atoms were then assigned atom map numbers based on the mapped neighbors they are bonded to, ensuring consistent tracking of reactive hydrogens across reactants and products. The pipeline proceeds by validating reaction syntax, generating reaction hashes, and deduplicating equivalent reactions to produce a clean, nonredundant set of full reactions (EVODEX-R).

### Phase 2: Partial Reaction Enumeration and Formula Pruning

Each full reaction (EVODEX-R) was decomposed into one or more partial reactions (EVODEX-P) using a combinatorial enumeration procedure implemented in split_reaction. This algorithm generates all possible nonempty reactant-product subcombinations and filters them to retain only those where at least one atom map number is shared between sides, indicating shared reactivity. Resulting SMIRKS strings were hashed to define unique EVODEX-P identifiers, with duplicates merged. Each EVODEX-P was abstracted to a formula-difference representation (EVODEX-F) using calculate_formula_diff(), which computes the net change in atom counts across the reaction and hashes this difference to create a grouping identifier. These formula diffs serve as a high-level abstraction of chemical change. To reduce noise, only EVODEX-F entries supported by five or more distinct EVODEX-Ps were retained, with the rest discarded alongside their corresponding EVODEX-Ps. This pruning step eliminated rare or malformed patterns, reducing downstream complexity and focusing the dataset on frequently observed, generalizable operators.

### Phase 3: Calculation and trimming of EVODEX-E

In Phase 3, we calculate all EVODEX-E operators from the retained EVODEX-P reactions and then prune them through a multi-stage filtering process. EVODEX-E represents the most specific operator family, capturing the full bonding and electronic environment involved in the transformation; because all other operator types are abstractions of E, we compute and prune E first to constrain the entire operator space.

All E operators are initially generated in their astatine form and deduplicated, linking each to their source EVODEX-P reactions. In Phase 3a, we discard any operators supported by fewer than ten partial reactions and then prune each set of sources down to five representative examples to reduce redundancy. In Phase 3b, we perform dominance pruning within each EVODEX-F group to remove structurally redundant E operators in favor of simpler alternatives. This step addresses the common issue of over-extended operators in conjugated systems, where extra atoms may be unnecessarily included in the reaction scope. The pruning algorithm ranks operators by increasing substrate atom count and eliminates any operator whose transformation can be reproduced by a simpler one. This projection-based dominance test ensures that only the most minimal yet predictive SMIRKS are retained. In Phase 3c, stable EVODEX.1 IDs are assigned across all operator types, completing the construction of the final EVODEX-E, -P, -R, and -F datasets.

### Phase 4: Enumeration of additional operators

The remaining operator families (C, N, Cm, Nm, Em) were derived from the retained EVODEX-P reactions by applying variant extraction settings to the same underlying SMIRKS. Each operator shares the same underlying reaction SMIRKS but differs in abstraction level based on the Boolean flags used in extract_operator. For each configuration, operators were extracted, converted from astatine placeholders to explicit hydrogens following hashing and consolidation.

### Phase 5: Calculation of EVODEX-M

After structural operator variants are computed, the pipeline derives formula-based mass information for analytical applications. Each EVODEX-P entry generates one EVODEX-F and one EVODEX-M, where the latter corresponds to the exact mass of the formula difference. This mass is calculated by multiplying each atom’s count by its precise atomic mass and summing the result. The full EVODEX-M table lists all such masses derived from the retained EVODEX-F entries. We also generate a filtered table, EVODEX-M_mass_spec_subset, containing only those masses corresponding to partial reactions where both the substrate and the product are single molecules. This one-to-one requirement ensures that the transformation reflects a clean mapping between a single substrate peak and a single product peak in a mass spectrum. It excludes scenarios such as fragmentations (1→n) or ligations (n→1), which are harder to interpret in isolation and may involve bespoke components. This stricter filter removes intramolecular and multi-species reactions, but a commented line in the code allows users to re-enable the broader set, including reactions where only one side is singular, if desired.

### Phase 6: Synthesis Subset

The pipeline also calculates a subset of the EVODEX-E operators specifically for product prediction applications such as biosynthetic pathway synthesis. To construct this subset, we first filter EVODEX-P reactions to exclude any containing ‘ubiquitous metabolites’—a predefined list of common cofactors, inorganic species, amino acids, and nucleotides presumed to already exist in cellular environments. We then identify the EVODEX-E operators associated with the remaining EVODEX-P entries and retain only those that are defined by exactly one substrate and one product. This excludes ligation, fragmentation, and other multi-species reactions involving bespoke metabolites. The result is a compact and biologically relevant synthesis subset. By the end of Phase 6, all retained operators are written as CSV files with astatines converted back to hydrogens.

### Phase 7: Website Generation

Finally, the curated data is transformed into a static website suitable for browsing. It operates on the final CSV files and generates a page for each operator of each type along with index pages containing a listing of all operators by type including the mass spectrometry and synthesis subsets. It also computes hyperlinks between related entries and includes adjacent information such as the source reactions. The pages also include SVG visualizations for each reaction or operator. The EVODEX-P pages consolidate all the information related to a partial reaction, including its EVODEX-R sources and derived operators. The generated website directory was uploaded to a separate GitHub repository^2^ and is served via GitHub Pages for public access.

## Results

### The Abstractions

The EVODEX project defines a family of abstractions to represent enzymatic reactions with chemical precision, capturing the underlying mechanistic transformations. These abstractions are instantiated in EVODEX.1 across ten operator types. Each abstraction reflects a distinct structural scope, as detailed below.

1. EVODEX-R (Full Reactions): An EVODEX-R represents an observed reaction that has been standardized and atom-mapped, with enzymes and unmodified chemicals removed. They are full reactions, chemically balanced, and include mapped hydrogens. The process of generating EVODEX-R includes the validation of reactions for parsability and stereochemical completeness.
2. EVODEX-P (Partial Reactions): EVODEX-Ps are derived from EVODEX-Rs by considering all possible combinations of substrates and products that share atom mappings. Non-matching atoms are removed, and the atom maps are reset to zero, resulting in a simplified SMILES string with unmapped atoms. These include singular substrate to singular product transformations, as well as n-to-n ones.
3. EVODEX-F (Formula Difference): This operator captures the net gain or loss of atoms during a transformation by calculating the atomic stoichiometry balance between substrates and products of an EVODEX-P. The resulting formula difference provides the most basic abstraction of a partial reaction and is efficiently calculated. They are used extensively in EVODEX for prefiltering and accelerating matching algorithms.
4. EVODEX-M (Mass Difference): EVODEX-M calculates the net change in molecular mass associated with the transformation captured by EVODEX-F. This operator is used in mass spectrometry applications, where this mass difference directly relates the masses of substrate and product ions.
5. EVODEX-Cm, -Nm, -Em (Matched Core, Nearest-Neighbor, and Electronic Operators): These operators are SMIRKS expressions with explicit hydrogens, containing only mapped atoms. EVODEX-Cm captures atoms with direct bond changes, EVODEX-Nm adds sigma-bonded neighbors, and EVODEX-Em includes all atoms sharing molecular orbitals with the core. Em operators most closely reflect classical arrow-pushing and are suited for classification and mechanism labeling.
6. EVODEX-C, -N, -E (Complete Operators): These are the full counterparts of the matched operators, including both mapped and unmapped atoms. They are used for product prediction, as they preserve the atom imbalance of partial reactions and enable reconstruction of plausible products.

### Synthesis Demonstration

To demonstrate a practical application of EVODEX operators, we developed a minimal synthesis projection script that applies EVODEX-E operators to candidate substrates. The central function, project_synthesis_operators, implements a greedy algorithm that applies each operator in the synthesis subset to a given molecule using RDKit. Although the approach is unfiltered and lacks performance optimizations such as substructure screening or fingerprinting, the modest number of operators and their typical single-substrate form keep runtimes manageable. This function is not intended to model enzyme specificity but instead to produce a comprehensive superset of mechanistically plausible one-step enzymatic products from a given substrate. It serves as a straightforward demonstration of how EVODEX abstractions can be used in forward prediction tasks.

We evaluated the synthesis subset using an interactive Colab notebook^3^ (also available in the repository as EVODEX_Synthesis_Demo.ipynb under notebooks), applying all EVODEX-E (synthesis) operators to a single input compound, n-propanol. The projected transformations included many common biochemical modifications: phosphorylation, acylation with various carboxylic acids, glycosylation with multiple sugars, sulfonylation, and O-alkylation. These reactions are consistent with known enzymatic activities across EC classes, suggesting that the subset captures the mechanistic diversity of biological chemistry. The output also included a smaller number of transformations that appear chemically implausible based on the static substrate structure, such as methylene C-methylation or hydroxylation of unactivated methylenes. For example, EVODEX.1-E677 corresponds to methylene C-methylation and enters the set through multiple reactions, including EVODEX.1-P1586, where the methylene lies between two carbonyls (e.g., a beta-diketone or malonyl-thioester analog), and EVODEX.1-P3059, where the methylated carbon is part of a CCN motif. Similarly, EVODEX.1-P534 maps a 16-carbon linear alkane to its terminal alcohol, producing a methyl-to-methanol operator, while EVODEX.1-P266 maps an internal methylene of a fatty acid to a secondary alcohol, yielding a methylene-to-alcohol operator. In both cases, the abstraction captures a plausible net transformation but omits the specific activation or intermediate states required for the actual chemistry. These examples highlight a key limitation: the operator abstraction does not capture transient mechanistic states that arise through cofactor interactions, conformational strain, or enzyme microenvironment. As a result, some predicted reactions may seem mechanistically implausible when viewed in isolation, even though they are derived from valid biochemical precedents. Figure 4 shows a static rendering of the full prediction set for n-propanol.

**Figure 4.**
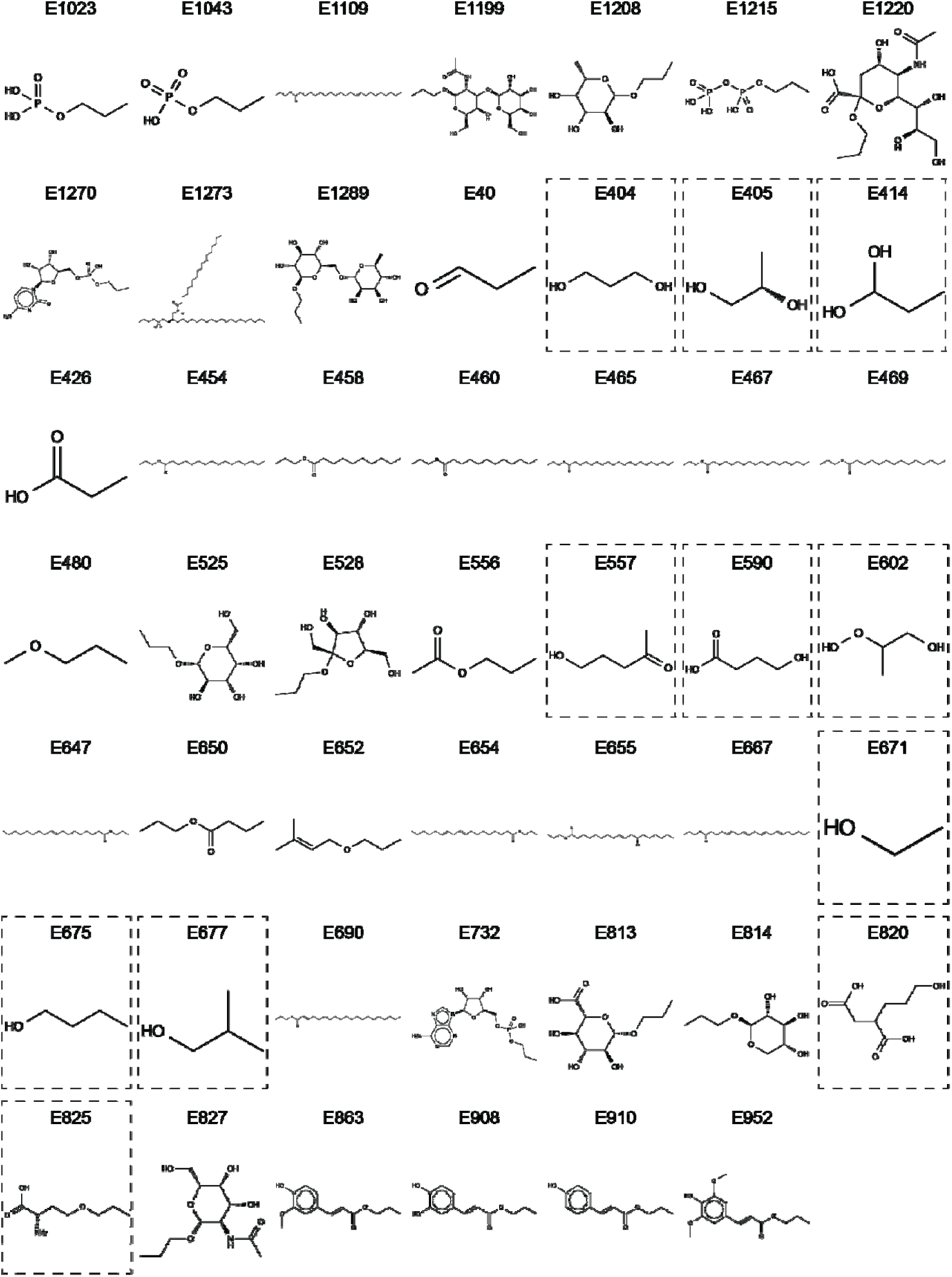
Predicted enzymatic products from n-propanol. Shown are the 48 unique products generated by projecting EVODEX-E (synthesis subset) operators onto n-propanol. Products outlined with a dashed rectangle represent mechanistically questionable predictions resulting from underspecified operators that lack full reaction context.

### Mass Spectrometry Demonstration

The mass spectrometry workflow is implemented in a dedicated notebook^4^ based on the EVODEX-M subset, a curated list of exact mass differences derived from substrate– product pairs in partial reactions. Each entry represents the net atomic mass change of a mechanistically plausible enzymatic transformation and is linked to one or more EVODEX-E operators. Given a known substrate and observed mass shift, the workflow identifies matching EVODEX-M entries within a specified tolerance and retrieves the corresponding EVODEX-E operators, which are then projected onto the substrate to generate candidate products. In a test case involving a steroid substrate with a mass shift consistent with methylation, the system returned both O- and C-methylated products. As discussed in the previous section, the C-methylations are likely implausible in isolation, but their inclusion reflects the design goal of returning a superset of structurally and mechanistically valid transformations. This shows how EVODEX enables chemically grounded interpretation of mass shifts.

### Evaluation Demonstration

To test operator-based plausibility scoring, we developed a two-stage evaluation module for assessing the mechanistic plausibility of enzymatic transformations. This functionality is particularly valuable when extracting reaction pairs from natural language sources, where automated parsing often results in hallucinations, mismatches, or misassignments.^20^ By checking whether a proposed substrate–product pair aligns with known mechanistic patterns, the EVODEX evaluation module offers a chemoinformatic filter to support reaction validation. Given a reaction and a target level of abstraction (EVODEX-F, -C, -N, or -E), the tool determines whether a corresponding operator exists that can plausibly explain the transformation.

The evaluation process consists of two stages. The first, assign_evodex_F, computes the formula difference between the substrate and product and checks whether this difference is represented in the EVODEX-F set. This serves as a fast plausibility filter based on atomic composition alone. If a match is found, the second stage, match_operators, retrieves all EVODEX operators of the specified type associated with that formula difference. Each operator is projected onto the substrate, and the resulting products are compared against the expected outcome. If a match is found, the operator is returned as a valid mechanistic explanation. This two-step strategy improves computational efficiency while maintaining accuracy.

To test this system^5^, we assembled a small set of real substrate–product pairs, including primary alcohol oxidations and amino acid transformations, and then generated decoy pairs by scrambling the inputs and outputs. Across all tested abstraction levels, the evaluation module correctly identified the true reactions and rejected all scrambled pairs. This result confirms that the operators can function as discriminative validators of mechanistic plausibility, providing a useful downstream tool for text mining, reaction curation, or quality control of biochemical databases.

### Pipeline outcomes and operator yields

We began with 349,458 atom-mapped reactions from EnzymeMap, all of which were successfully loaded using RDKit and passed initial validation for atom balance and SMILES syntax. Each was then converted to an astatine-labeled form to facilitate improved handling of reactive hydrogens during atom mapping. A total of 1,060 reactions (0.3%) were lost during this remapping step due to processing errors. After deduplication, we retained 36,868 unique full reactions (EVODEX-R), representing roughly a 10-fold reduction, consistent with the observed mean of 9.45 source reactions per EVODEX-R. This redundancy reflects the enzyme- and organism-specific nature of the original annotations, where orthologous transformations are represented repeatedly across taxa.

Phase 2 expanded the dataset to 186,258 EVODEX-P partial reactions, without additional loss. The increase reflects the combinatorial decomposition of each full reaction into multiple subtransformations, while overlap among common motifs (e.g., NAD⁺/NADH) led to some consolidation. A large fraction of the resulting EVODEX-Ps are unique to a single full reaction and reflect limited atom overlap, such as pairings that share only a hydrogen. This produces a broad distribution that includes many rare, idiosyncratic reactions. From these partial reactions, 29,832 unique EVODEX-F formula differences were computed, with 10 discarded due to processing errors. Applying a support threshold of five or more source EVODEX-Ps yielded a pruned set of 3,352 formula classes, eliminating approximately 89% of the EVODEX-Ps and discarding rarely observed or poorly generalizable patterns.

Of the remaining partial reactions, 97% were successfully interpreted as EVODEX-E operators, producing 35,978 distinct transformation patterns. To focus the dataset on well-supported and interpretable reactions, we retained only EVODEX-E operators backed by at least 10 EVODEX-Ps, an empirically chosen threshold that removed over 90% of candidates while substantially improving signal-to-noise. Operators with lower support were more likely to contain mapping artifacts or represent highly specific transformations. An additional 290 entries were lost during conversion from astatine back to hydrogen, and 107 were pruned as structurally redundant during dominance filtering, leaving a final curated set of 1,404 EVODEX-E operators.

All remaining operator families (C, N, Em, Cm, Nm), along with mass-difference entries (EVODEX-M) and a synthesis-focused subset, were deterministically generated from the retained EVODEX-Ps without further loss. These collectively form the complete EVODEX.1 operator release, which serves as the foundation for downstream applications in synthesis planning, mechanistic modeling, and mass spectrometry interpretation. Table 1 summarizes the major phases and runtimes of the pipeline. Final operator counts are shown in Table 2. A full breakdown of intermediate statistics is available in the EVODEX GitHub repository as pipeline_results.csv.

**Table 1.** Overview of EVODEX Pipeline Phases and Runtime.

| Phase | Description | Runtime (s) |
| --- | --- | --- |
| 1 | Reaction validation, deduplication, atom mapping | 7477.29 |
| 2 | Partial reaction generation and formula pruning | 3068.62 |
| 3 | EVODEX-E computation | 1464.73 |
| 3a | Source support filtering and operator trimming | 4.67 |
| 3b | Dominance pruning | 122.59 |
| 3c | Publishing and ID assignment | 44.34 |
| 4 | Generation of C, N, Em, Cm, Nm operators | 414.92 |
| 5 | EVODEX-M mass delta generation | 0.39 |
| 6 | Synthesis subset selection | 15.20 |
| 7 | Website and documentation build | 237.64 |

**Table 2.**
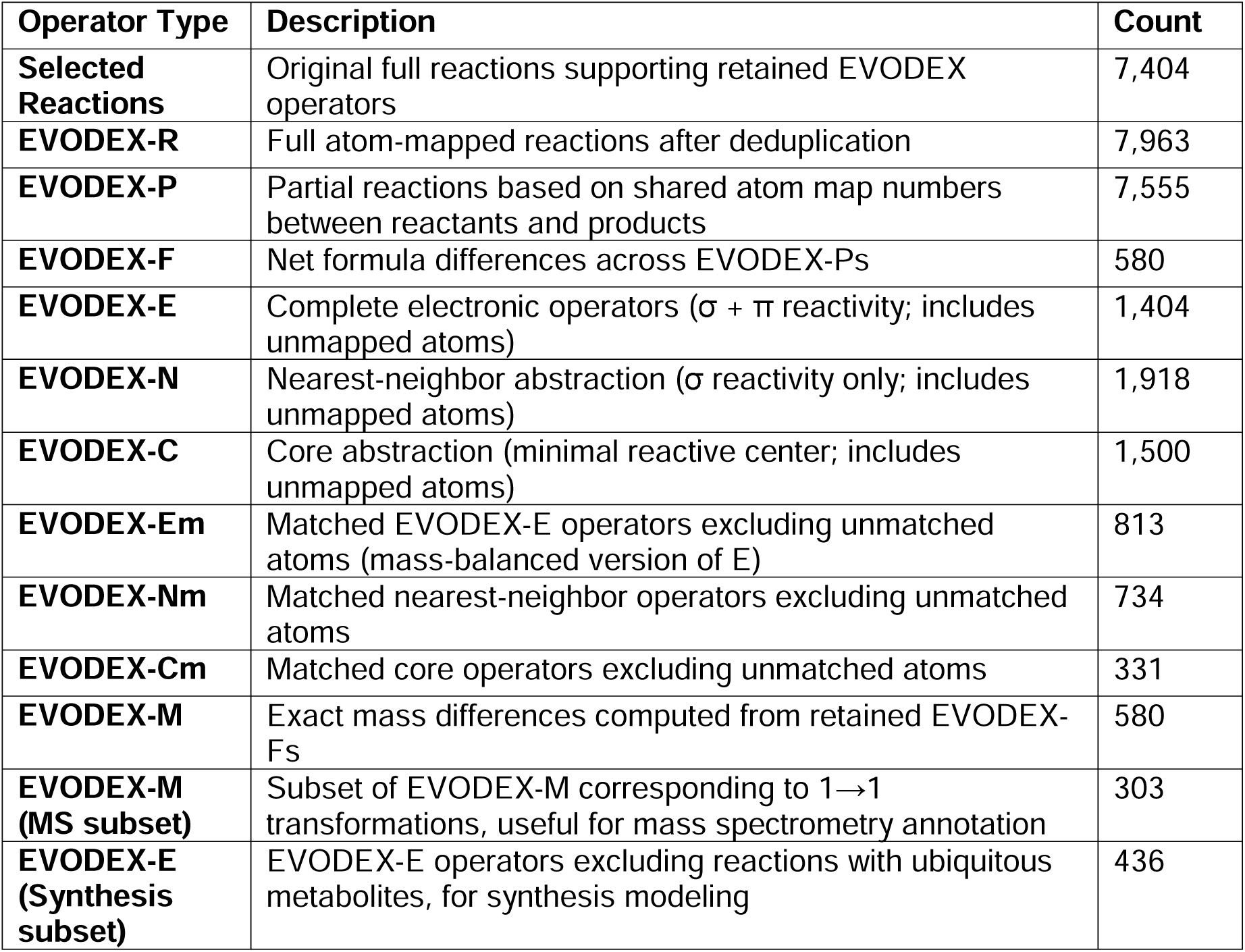
EVODEX.1 Operator Types and Final Counts.

### Functional Coverage of Naïve Reactions

To evaluate the coverage of EVODEX.1, we assembled a test set of 1000 enzymatic reactions randomly sampled across EC classes using ec_coverage_sampler.py. Reactions were stripped of ubiquitous metabolites (decofactor.py) and reduced to single substrate–product pairs, represented as SMILES strings without atom maps or hydrogens. This filtered set was passed to ec_coverage_evaluation.py, which applied assign_evodex_F to compute formula differences and match_operators to identify matching EVODEX-E operators. All 1000 reactions were processed successfully, with no evaluation failures.

Of the total, 906 reactions (91%) had a matching EVODEX-F formula difference, indicating stoichiometric consistency with known partial transformations. At the EVODEX-E level, 618 reactions (62%) returned at least one matching operator, with 100 reactions matching multiple operators. This likely reflects residual structural redundancy that were not fully eliminated during the dominance pruning step in mining. The most frequent individual E operator (EVODEX.1-E41, alcohol to ketone) matched 74 reactions, and overall hit distribution was weighted toward well-supported transformations. The full evaluation report and per-reaction results are included in analysis_reports/evaluation_stats.txt.

### Source Coverage by EC number

The observed 91% formula-level match rate in the functional evaluation prompted a closer examination of whether EVODEX-E operators were broadly representative or simply overfit to a redundant input set. To investigate this, we assessed the enzymatic diversity of the reactions retained during mining by comparing their EC numbers to those in the full EnzymeMap dataset. This analysis, performed using ec_comparison.py, compared raw_reactions.csv (all input reactions) against selected_reactions.csv (those supporting final EVODEX-E operators). Results are provided in ec_coverage_report.csv in analysis_reports and summarized in Table 4.

**Table 3.** Reaction-level outcomes from evaluating 1000 test reactions using EVODEX-F and EVODEX-E.

| Metric | Value |
| --- | --- |
| Reactions evaluated | 1000 |
| Reactions with EVODEX-F match | 906 |
| Reactions with EVODEX-E match | 618 |
| Reactions with multiple EVODEX-E matches | 100 |
| Most frequent E operator (EVODEX.1-E41) | 74 hits |
| Unique E operators matched ($\geq 1$ hit) | 117 |
| Reactions with no E operator match | 382 |
| Average evaluation time per reaction | 0.0354 |

**Table 4.** EC Coverage Summary. Comparison of EC numbers in the full EnzymeMap dataset and those retained as support for EVODEX-E operator generation.

| EC Level | Selected Count | Raw Count | Overlap Count | Coverage (%) |
| --- | --- | --- | --- | --- |
| 1 | 6 | 6 | 6 | 100 |
| 2 | 48 | 61 | 48 | 78.7 |
| 3 | 135 | 204 | 135 | 66.2 |
| 4 | 1770 | 4552 | 1770 | 38.9 |

At EC level 1, all six major enzyme classes are represented in the retained set. However, coverage declines with increasing classification specificity, dropping to 79% at level 2, 66% at level 3, and 39% at level 4. Since only a minority of specific EC entries were retained, the results show that the final operator set was not overfit to the full dataset. Instead, EVODEX abstractions are supported by a compressed subset of reactions that recapitulate a much larger space of mechanistic content.

### Analysis of omitted reactions

To assess the full diversity of chemistries in the dataset, we ran the EVODEX-R reactions through extract_operator without applying pruning for formula frequency or operator dominance. This unfiltered run over 36,868 reactions yielded 4,419 unique EVODEX-E operators, with only 243 processing failures. This defines an upper bound on the reaction space. As shown in Figure 5A, the distribution of support per operator is sharply skewed: over half (56 percent) are singletons, supported by only one reaction, with a dramatic log-scale dropoff. These unicorn operators arise from various causes, shown in Figure 5B. Some are valid but rare (e.g., extensive delocalization or unusual elements like Cr), some reflect stereochemical complexity, and others result from mapping errors in the source data or artifacts from astatine substitution. Because EVODEX-E extraction depends entirely on atom maps, poor mappings can produce incorrect or overly specific operators. However, projection of mined operators during synthesis prediction is mapping-independent and based only on structure, so valid operators from correctly mapped reactions can still recover the correct transformation. Although these low-support cases were excluded from the final release to improve robustness, the curated set still explains 91% of tested reactions at the formula level and 62% at the mechanistic level. These levels of coverage likely result from both general operators explaining complex reactions and the resilience of structure-based projection. While the approach cannot capture every edge case in a general-purpose dataset, organism-specific sets can be completed through EVODEX assignment and manual curation.

**Figure 5.**
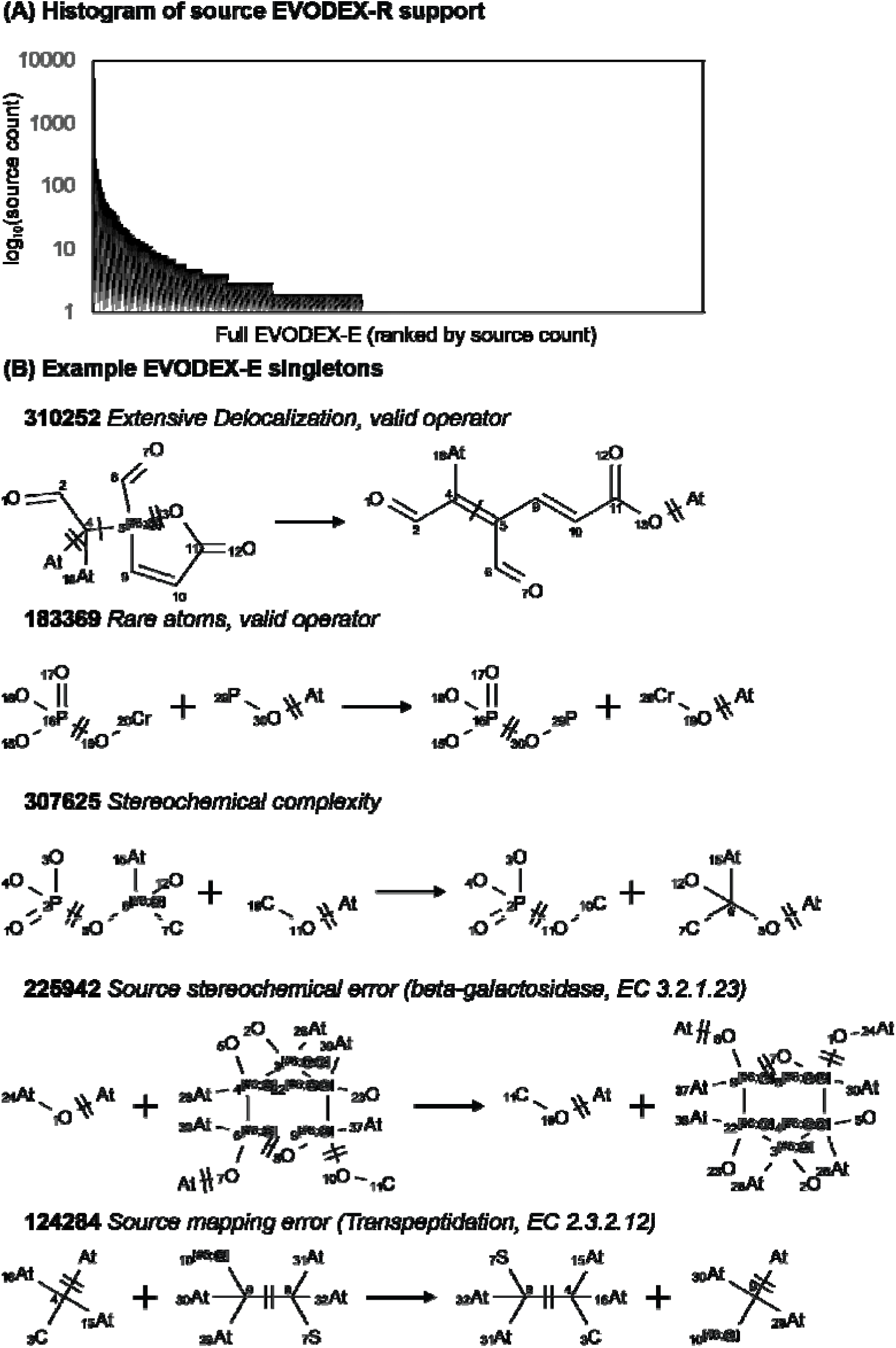
Source support and diversity of EVODEX-E operators. (A) Histogram of EVODEX-R reactions per EVODEX-E operator, showing a skewed distribution with many low-support cases. (B) Examples of singleton EVODEX-E operators arising from valid but rare chemistry (e.g., delocalization, uncommon elements, stereochemical complexity) or likely artifacts (e.g., mapping and stereochemical errors). Operators with low source support were excluded from the final release set.

### Manual Review of Edge Cases

We manually reviewed the first 50 operators in the synthesis subset in detail, including those with unusual or atypical chemistries, and found no chemically invalid cases. Some operators appear under-specified, representing transformations like C-methylation or hydroxylation at unactivated positions, where the true mechanism likely involves features not captured in the abstraction. A few also retain irrelevant stereochemical labels. Spot-checks of the other operator families revealed minor issues, such as extraneous atoms in some C operators due to an unknown bug. Overall, the dataset has been refined through multiple iterations, and the final operators appear chemically sound and mechanistically plausible within their intended scope.

## Discussion

### The Need for a Minimal, Objective Abstraction

The Electronic Reaction Operator (ERO) defines an abstraction of enzymatic reactivity that is both minimal and auditable. In contrast to curated rules or machine-learned predictions, EROs are derived directly from molecular structure and atom mapping using a deterministic algorithm. This ensures that each operator corresponds to a real, interpretable transformation—one that a chemist could reconstruct from first principles. In this sense, EVODEX provides a language for chemical mechanism that is accessible both to humans and software, independent of statistical inference.

### The Physicoorganic Basis for the ERO

Reactivity in organic chemistry is governed by the configuration of molecular orbitals, specifically the sigma framework and any adjacent pi systems involved in conjugation or electron delocalization. EROs are built on this foundation, including not only atoms undergoing bond changes but also neighboring atoms whose orbitals are essential to the mechanism. This distinguishes EROs from more abstract operator types such as EVODEX-C and EVODEX-N, which include only the core atoms or their immediate sigma neighbors. These coarser abstractions often collapse mechanistically distinct transformations. For instance, a primary alcohol and a carboxylic acid are nearly indistinguishable at the C level and only modestly differentiated at N. An α,β-unsaturated carboxylate, such as an acrylate, shares the same core atoms but undergoes entirely different chemistry due to extended conjugation. As shown in Figure 6 and the corresponding notebook, the operator, mined from a Michael addition of thiomethane to acrylate, is projected onto a panel of possible substrates. At the C level, any terminal alkene is predicted to react, ignoring the role of the carbonyl; at the N level, this error persists.

**Figure 6.**
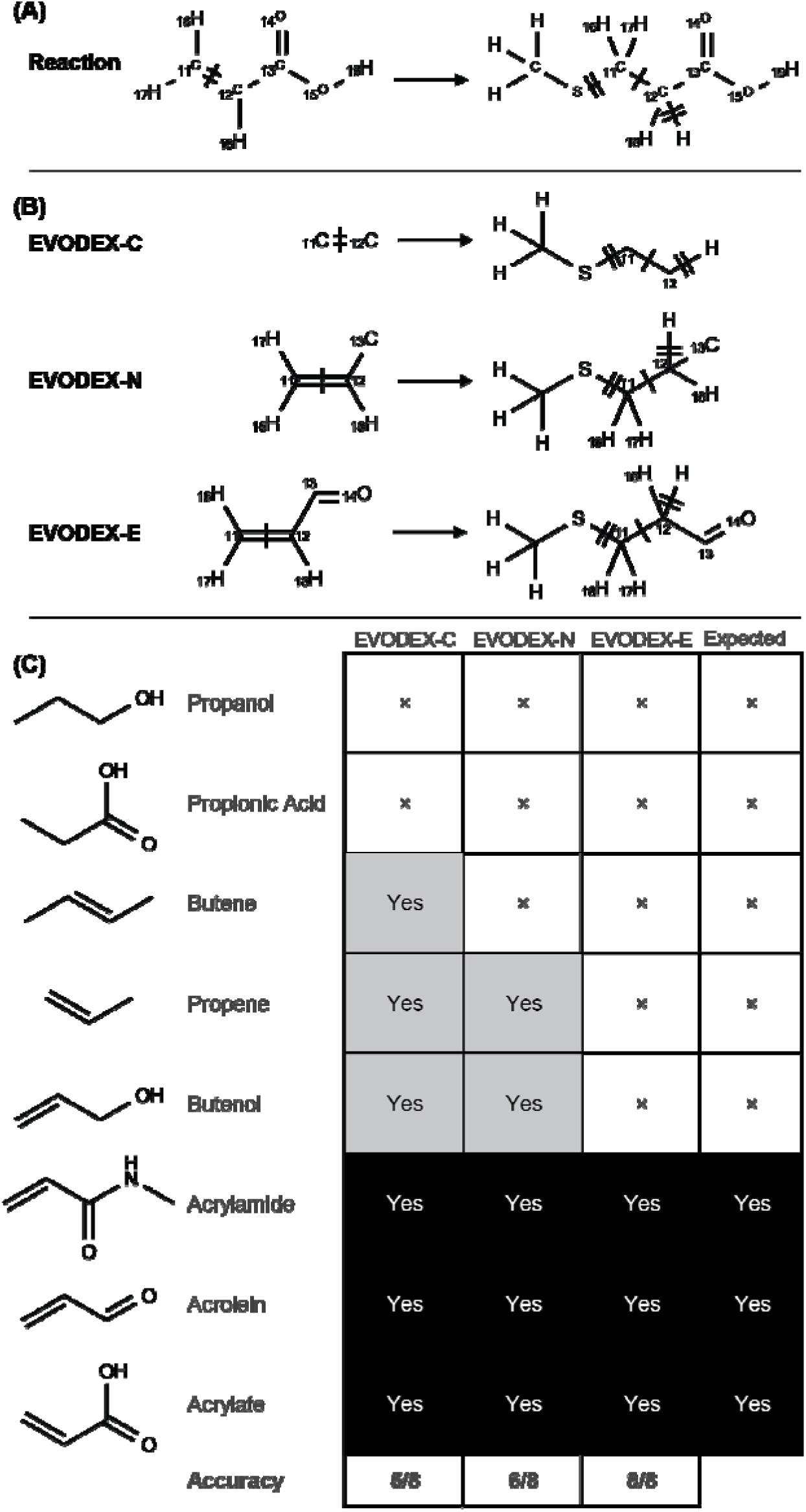
Effect of abstraction level on reactivity prediction for a Michael addition. (A) Source reaction: conjugate addition of thiomethane to acrylate, used to mine the EVODEX operator. (B) Substrate panel: a series of analogs with varying degrees of conjugation and electron-withdrawing character. (C) Predicted reactivity at three abstraction levels: Core (C), Nearest-Neighbor (N), and Electronic (E).

### Limitations of the ERO Abstraction

While the Electronic Reaction Operator is constrained by the granularity of available data. In principle, an ERO would include all atoms contributing to bonding at any point along the reaction coordinate, including intermediates and transition states. In practice, biochemical datasets provide only atom-mapped substrates and products, limiting the analysis to endpoints. As a result, transiently involved atoms may be excluded. For example, in malonate decarboxylation, the central carbon forms an enolate intermediate conjugated with two carbonyls, but this configuration is absent in both substrate and product. The resulting operator omits key atoms and is mechanistically incomplete, a frequent cause of implausible predictions in synthesis tasks. Conversely, some operators include atoms not directly involved in the transformation, such as conjugated systems or stereocenters preserved from non-stereospecific reactions. To address this, we apply dominance pruning to discard operators that are strict superstructures of simpler, equally supported variants.

### Expanding the Operator Space

While EVODEX-E operators must be mined from data due to their dependence on real structural and electronic contexts, the matched Cm operators offer an opportunity for first-principles enumeration. Because Cm operators describe only directly reacting atoms with explicit hydrogens, they collapse most transformations into a small vocabulary of local patterns—typically of the form X–Y to X–Z involving common biochemical elements like C, H, N, O, P, and S. This boundedness makes them suitable for full enumeration: the space of chemically reasonable reacting fragments is small and tractable. This contrasts with E operators, where extended conjugation and context-sensitive orbital interactions introduce unbounded variability. Thus, Cm forms a finite, enumerable space ideal for reaction classification and ML features, while E defines the open, mechanistically complete space required for synthesis and reactivity modeling.

### Defining Partial Reactions

Partial reactions are a core abstraction in EVODEX, enabling the modeling of enzymatic chemistry even when full reaction data is unavailable. Many reactions in biochemical databases are incomplete, ambiguously mapped, or involve multiple substrates and cofactors whose roles are context-dependent. By isolating only the atom-mapped substrate-product pairs that share mapped atoms, EVODEX constructs partial reaction operators that describe a single mechanistic transformation within a broader enzymatic context. This introduces complexities such as unbalanced reactions, unmapped fragments, and the need to handle hydrogen atoms explicitly. However, it avoids brittle assumptions about enzyme context, reaction balancing, or cofactor roles, and allows mechanistic reasoning over chemically participating fragments. This enables operator mining at scale from imperfect data, supporting tasks like synthesis planning and mass spectrometry interpretation without requiring full pathway annotation or complete reaction context.

### Rethinking Cofactors in Computational Models

A key step in defining partial reactions is deciding which molecules to exclude as cofactors. While this may seem straightforward, it turns out to be a nuanced problem. Traditionally, cofactors are defined as non-protein molecules that assist enzymes, but this identity-based definition breaks down in large-scale biochemical datasets. Molecules like ATP or NADH act as cofactors in some contexts and as substrates in others, making static exclusion lists unreliable. EVODEX avoids this ambiguity by applying a structural rule: a molecule is only included in a partial reaction if it contains atoms that are atom-mapped to a counterpart on the opposite side of the reaction, ensuring objective inclusion based on direct chemical participation. For synthesis tasks, EVODEX introduces a curated set of ubiquitous metabolites, which form a broader category than classical cofactors and include media components, core dogma players, and common donors. This allows reactions involving ubiquitous metabolites, such as NADH plus alcohol yielding an aldehyde, to be simplified to alcohol yielding aldehyde, since the cofactor is presumed present. In contrast, reactions involving non-native compounds, like a hypothetical synthetic nucleotide sugar dGTP-F-glc plus alcohol yielding alcohol-F-glc, are not reducible.

### Limits of Mapping Accuracy

Atom-to-atom mapping is a physically real and in principle auditable property, one that can be validated using isotope labeling and mechanistic studies. However, high-quality mappings are rare. Most biochemical datasets rely on heuristic, ML, or rule-based methods, which can produce systematic errors, and few assign maps to hydrogens. We identified multiple mis-mapped reactions in the source data (see Figure 5). These caused incorrect or misleading operators, especially when mappings were accepted with few supporting reactions. We mitigated this by requiring at least ten supporting examples per operator, which excluded many artifacts that had passed a five-example threshold. RXNMapper^21^ was evaluated for an alternate set of mappings. Despite its high reported accuracy^22^ the error rate was higher than the frequency of rare but legitimate transformations.

### Hydrogen Handling and the Astatine Hack

Hydrogen atoms must be treated explicitly to represent partial reaction operators faithfully. Many enzymatic reactions, like transamination, involve mechanisms that do not formally involve redox, but appear as redox transformations when viewed in isolation from the full reaction context. This apparent change in oxidation state arises from hydrogen shifts that are invisible when hydrogens are implicit. The effects are demonstrated in the RDKit_RO_Projection_Hydrogen_Study.ipynb notebook in the GitHub repository. Most cheminformatics libraries omit or inconsistently handle hydrogens, which breaks this accounting. Of the toolkits tested (ChemAxon, Indigo, and RDKit) only ChemAxon consistently preserved explicit hydrogens across all required operations. Due to licensing constraints, we adopted RDKit, but substituted hydrogen atoms with Astatine, an unused element that preserves valence behavior. This workaround made the algorithm tractable but introduced stereochemical artifacts, data loss, and significant complexity. Fortunately, RDKit supports explicit hydrogens during operator projection, allowing the final EVODEX operators to be presented without Astatine substitution.

### Cheminformatics Language and Toolkit Limitations

Current cheminformatics languages like SMARTS and SMIRKS describe molecules in terms of bond order rather than hybridization. This creates ambiguity around resonance and aromaticity, where multiple valid representations exist for the same underlying electronic structure. The problem is especially acute in biomolecules with fused heterocycles, where differences in kekulé forms or aromatic flags can obstruct otherwise identical matches. Hybridization provides a more stable and physically meaningful description, collapsing these variants into a single representation. EVODEX handles this internally using hybridization-aware graphs, but this information cannot be expressed in current cheminformatics formats, limiting compatibility and precision.

## Conclusion

EVODEX provides a mechanistic foundation for modeling enzymatic reactivity by abstracting transformations into small, auditable units derived from structure and atom mapping. It does not model enzyme specificity or biological context but instead captures what is chemically possible, creating a stable base for downstream models to add constraints related to binding and biophysical context. This separation of mechanism from context allows EVODEX to support diverse applications including synthesis prediction, mass spectrometry interpretation, and reaction plausibility checks. By defining a compact, reusable set of operators from a large and noisy dataset, EVODEX demonstrates that enzymatic reactivity can be systematically represented, enabling interpretable and scalable reasoning in computational biochemistry.

## Authors’ Contributions

J.C.A. led the development, analysis, and writing of this work. Early versions of the code were implemented in Java using the Indigo toolkit by H.L., D.T., and H.Z. The project was later transitioned to Python and RDKit, with contributions from L.L.C., who also focused on evaluating input biochemical datasets and mapping tools. G.X. and C.Z. contributed to the operator matching algorithm used to assign operators to reactions. All authors contributed to the development of the software and manuscript text. The authors used OpenAI’s ChatGPT throughout the project for content discussion, code generation, manuscript drafting, and editing.

## Acknowledgements

EVODEX builds on ideas first explored during the Act Synthesizer project at UC Berkeley, supported indirectly by DARPA Living Foundries. We thank Saurabh Srivastava, Ras Bodik, and Sanjit Seshia for their contributions to early conceptual development. Related work at 20^n^ and insights from DARPA’s Living Foundries: 1000 Molecules program helped shape the ideas behind EVODEX, though the project itself was not directly funded.

## Supplemental Information

The source code^1^ and browsable website^2^ are available on Github. Additional data from analysis of the mining process is available in the EVODEX repo^1^ under analysis_reports.

